# Latent Interacting Variable-Effects Modeling of Gut Microbiome Multi-Omics in Inflammatory Bowel Disease

**DOI:** 10.1101/2022.07.08.499280

**Authors:** Javier. E. Munoz, Douglas. K. Brubaker

## Abstract

Latent Interacting Variable Effects (LIVE) modeling is a framework to integrate different types of microbiome multi-omics data by combining latent variables from single-omic models into a structured meta-model to determine discriminative, interacting multi-omics features driving disease status. We implemented and tested LIVE modeling in publicly available metagenomics and metabolomics datasets from Crohn’s Disease and Ulcerative Colitis patients. Here, LIVE modeling reduced the number of feature correlations from the original data set for CD and UC to tractable numbers and facilitated prioritization of biological associations between microbes, metabolites, enzymes and IBD status through the application of stringent thresholds on generated inferential statistics. We determined LIVE modeling confirmed previously reported IBD biomarkers and uncovered potentially novel disease mechanisms in IBD. LIVE modeling makes a distinct and complementary contribution to the current methods to integrate microbiome data to predict IBD status because of its flexibility to adapt to different types of microbiome multi-omics data, scalability for large and small cohort studies via reliance on latent variables and dimensionality reduction, and the intuitive interpretability of the linear meta-model integrating -omic data types. The results of LIVE modeling and the biological relationships can be represented in networks that connect local correlation structure of single omic data types with global community and omic structure in the latent variable VIP scores. This model arises as novel tool that allows researchers to be more selective about omic feature interaction without disrupting the structural correlation framework provided by sPLS-DA interaction effects modeling. It will lead to form testable hypothesis by identifying potential and unique interactions between metabolome and microbiome that must be considered for future studies.

**AUTHOR SUMMARY:** Latent Interacting Variable Effects (LIVE) modeling integrates microbiome multiomics features by encoding them in a set of latent variables (LVs) from single-omic sparse Partial Lease Squares models, and then combine these LVs into structured metamodel to determine the most discriminative features driving IBD. We used publicly available metagenomic and metabolomics data from Crohn’s Disease and Ulcerative Colitis patients to develop LIVE modeling. LIVE modeling reduced data dimensionality efficiently and identified statistical interactions among microbiome multi-omics data, which can be visualized as a mineable network data structure. LIVE modeling confirmed features previously reported and revealed novel microbiome interactions in IBD. LIVE offers a flexible framework for multi-omic modeling that may aid in interpretation of complex microbiome datasets.

## INTRODUCTION

Alterations in the composition and function of intestinal microbiota are a hallmark of Crohn’s Disease (CD) and Ulcerative Colitis (UC), two subtypes of Inflammatory Bowel Diseases (IBD) [1-4]. These alterations trigger local and systemic host inflammatory responses that progressively damage the intestines over time [5-6]. A hypothesis of how alterations in gut microbiota drive chronic inflammation in Crohn’s Disease and Ulcerative Colitis is through the gut metabolome which may act as an interface between the host and microbiome [7]. However, these mechanisms of metabolome-mediated host-microbiome interactions are poorly understood. A better understanding of how gut microbiome alterations drive IBDs could enable less-invasive diagnostic methods for CD and UC, facilitate identification of therapeutic biomarkers, or reveal strategies to prolong disease remission and reduce the likelihood of relapse [8]. As validated pathways of host-microbiome interactions in IBD continue to be established, interpretable computational models of microbiome multi-omics data could potentially reveal translatable insights into IBD progression and drug-resistance, data which is increasingly available from large cohorts of CD and UC patients.

Much of the current state of the art to determine microbe-metabolite associations is focused on pairwise correlation approaches, supervised machine learning, unsupervised clustering methods, and multivariate linear models [9-11]. These techniques are often challenged by the dimensionality of the multi-omics data, often do not account for a correlation structure within microbiomes or metabolomes and are challenged by high multiple hypothesis testing burdens. Some examples of how computational workflows try to address high data dimensionality include applying feature selection methods to random forest (RF) classifiers or reducing the feature space with latent variable methods like principal component analysis (PCA), and partial least squares – discriminant analysis (PLS-DA). Latent variable approaches provide additional advantages because of the global covariance-based frameworks that reduce data dimensionality by encoding the data on latent variables (LV) that weight the relative importance of the original features [13]. PLS-DA uses latent variables to establish a system of linear relationships between blocks of independent and dependent variables which estimate causal relationships between observed indicators and their latent variables.

A variant of PLS-DA, sparse PLS-DA (sPLS-DA), uses Lasso Penalization to select the most discriminant features to create the LVs to classify the data [16-18]. An example of these methods was presented by Priya *et al*., 2022 in which host transcriptomic and gut microbiome data were integrated in a machine learning framework that uses sparse canonical correlation analysis and Lasso penalize regression to characterize the most relevant associations between gut microbiota and host genes and pathways across three intestinal diseases [19]. This regularization penalty on PLS latent variables helps reduce the curse of dimensionality in high-dimensional data. Critically however, if multiple -omics data types are modeled on the same LVs, the Lasso penalty may be overly-restrictive and exclude consequential biological features. Therefore, there is a need to balance dimensionality reduction with preserving potentially mechanistic biological information in different molecular data types.

Here, we introduce a computational framework for causal modeling of microbiome multi-omics data based upon a Partial Least Square Path Modeling (PLS-PM) framework, and which will be denoted as Latent Interacting Variable-Effects (LIVE) Modeling. PLS-PM approaches incorporate the advantages of LV modeling for data dimensionality reduction while also including information about the modeled system through structural connections between model components, allowing the model to encode hypotheses of causality [14-15]. Here, we construct PLS-PM models of microbiome composition, metabolomics, and bacterial proteomics data from a cohort of 155 IBD patients and controls in which each molecular data stream is encoded as a set of latent variables. We then construct a structural model in which we quantify interactions between latent variables with linear model interaction effects. We show that LIVE Modeling accounts for global data covariance structure in the single-omic LV’s, efficiently reduces data dimensionality, and allows us to identify statistical interactions between microbiome multi-omics data associated with disease onset in UC and CD.

## RESULTS

### Model Description

LIVE modeling integrates multi-omics data while preserving covariance structure within data modalities. sPLS-DA models are built to model the relationship between individual types of molecular features and an output variable. A variety of microbiome multi-omics data types, such us relative abundances of genes, taxa, and metabolites, can be analyzed individually with respect to the dependent variable in their own dimensionless latent variable space. By building a *bottom-up* multi-omic model from individual grassroots models of single data types, this framework can incorporate several types of microbiome multi-omics data as inputs, determining the most discriminative latent variables per data type. Second, the latent variables are extracted from the single-omic sPLS-DA models and integrated together as main and interaction effects terms in a generalized linear model, formalizing a structured relationship between the response variable and multiple predictive -omic features. This allows us to evaluate data type-specific effects and determine how the predictive power of data types depend on each other. Using the LV’s as regression terms preserves internal covariance within data type LVs and simplifies the interpretability of the multi-omic effects in the model through analysis of the regression coefficients. Finally, the relationships between multi-omics data types and specific features implicated by the meta-model can be studied by modeling the relationships between microbes, metabolites, and enzymes via Spearman correlation analysis and PLS Variable Importance of Projection (VIP) scores (Figure 1).

**Figure 1.**
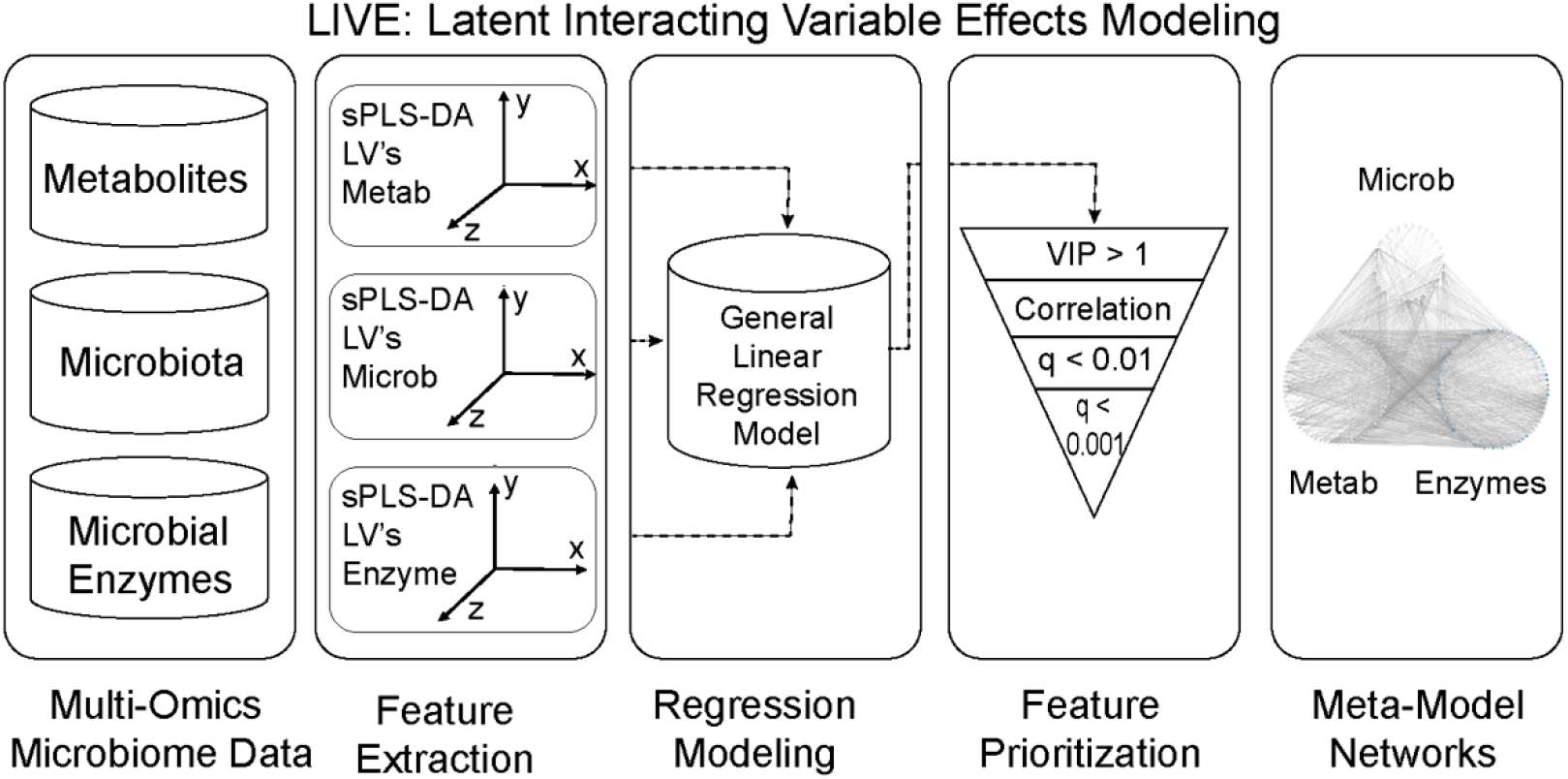
Latent Interacting Variable Effects (LIVE) Modeling Workflow. Multi-Omics Microbiome Data is encoded in a set of Sparse Partial Least Square Models. Then, structured model combines discriminative latent variable per each single-omics sPLS-DA models. Next, significant discriminative and predictive omic features are thresholding by VIP, correlational analysis, and q-values to build meta model networks.

We applied our LIVE modeling framework to predict CD or UC status versus control from publicly available metabolomics, and metagenomics data obtained from the PRISM cohort (the Prospective Registry in IBD Study at MGH) and reported by Franzosa *et al*., in 2019. In this study, gut metabolic profile and microbiome composition in IBD were characterized by analyzing stool samples of 155 patients diagnosed with Crohn’s Disease (68 patients), Ulcerative Colitis (53 patients), and non-IBD control (34 patients). The collected stool samples were subjected to shotgun metagenomic sequencing to determine taxonomic compositional and functional potential, as well as four liquid chromatography tandem mass spectrometry (LC-MS) methods to quantify composition of polar metabolites, lipids, free fatty acids, and bile acids. The datasets contained the relative abundances of 201 microbiota features, 3829 metabolic features, and 2113 microbial enzyme features [9]. To validate our model trained on PRISM data and determine performance metrics, we used a public independent validation cohort also reported by Franzosa *et al*., consisting of 22 control subjects from LifeLines-DEEP (LLD) general population study, and 43 IBD subjects from a study of the Department of Gastroenterology and Hepatology at University Medical Center Groningen (UMCG).

### Single-omic sPLS-DA are as predictive of Crohn’s Disease status as naive multi-omics models

Before combining single-omic latent variables (LV) from sparse Partial Least Square Discriminant Analysis (sPLS-DA) via the LIVE multi-omic meta-model, we assessed the disease-status predictive power of these data streams individually training on the PRISM cohort and testing on the LLD cohort for Crohn’s Disease and Ulcerative Colitis. The individual microbiome, metabolome and microbial enzyme sPLS-DA models were significantly predictive of CD status. (Metabolome AUC = 0.959: Figure 2a and 2b, Microbiome AUC = 0.972: Figure 2e and 2f, Enzymatic composition AUC = 0.939: Figure 2i and 2j). The most discriminative features per each single sPLS-DA models were sorted by the PLS Variable Importance of Projection (Figure 2c, 2g, 2k) and the top features are listed here: the most discriminative metabolites for CD are the flavonoids, steroidal glucuronide conjugates, fatty acids and conjugates, dipeptides, terpene glycosides, tetrapyrroles and derivates, benzenesulfonic acids and derivatives, triterpenoids, benzopyrans and alpha-acyloxy carbonyl compounds (Figure 2d). The most discriminant microbial species are *Coprococcus catus, Subdoligranulum unclassified, Alistipes shahii, Roseburia hominis, Bacteriodales bacterium* ph8, *Eubacterium ventriosum, Ruminococcus obeum, Gordonibacter pamelaeae* (Figure 2h). Finally, the most discrimative enzymatic features are 1.14.13.81: Magnesium-protoporphyrin IX monomethyl ester (oxidative) cyclase, 1.1.1.35: 3-hydroxyacyl-CoA dehydrogenase, 3.5.3.9: Allantoate deiminase, 3.1.4.1: Phosphodiesterase I, Saccharopine dehydrogenase (NAD(+), L-lysine-forming) (Figure 2l).

**Figure 2.**
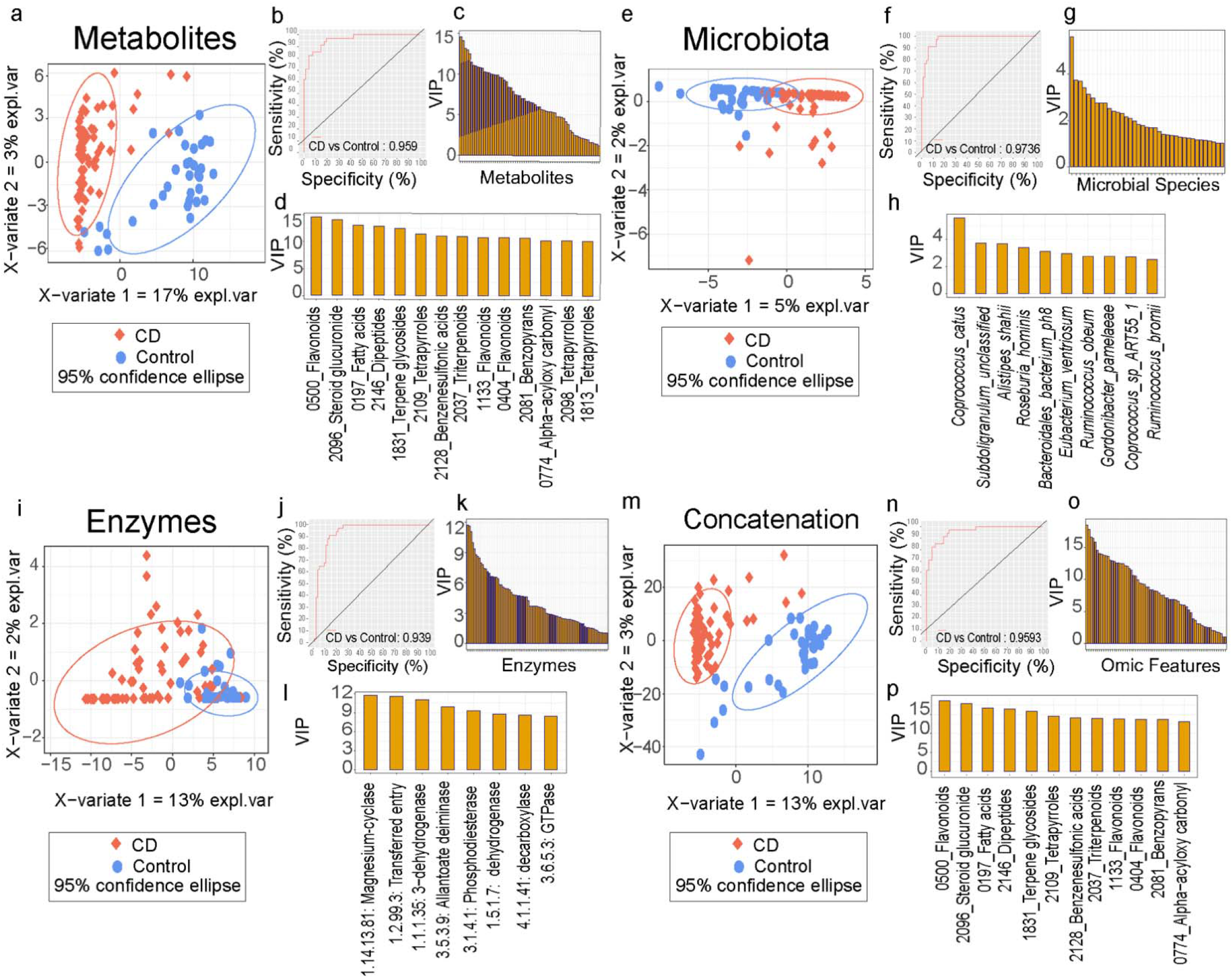
sPLS-DA model performance and variance importance of projections (VIP) for CD. (a) sPLS-DA sample scores plot for metabolomics data. (b) ROC curve for metabolomics sPLS-DA model. (c) VIP score histogram for metabolites. (d) Top 14 metabolites by VIP score. (e) sPLS-DA sample scores plot for microbiota composition data. (f) ROC curve for microbiota sPLS-DA model. (g) VIP scores histogram for microbiota. (h) Top 10 microbial taxa VIP scores. (i) sPLS-DA sample scores plot for enzymes data. (j) ROC curve for enzymes sPLS-DA model. (k) VIP score histogram for enzymes. (l) Top 8 enzymes by VIP score. (m) sPLS-DA sample scores plot for concatenation control multi-omic model. (n) ROC curve for concatenation control model. (o) VIP score histogram for concatenation control model. (p) Top 12 feature VIP scores for the control model.

As a computational control, we trained sPLS-DA models on a concatenated matrix that integrated all data types (microbiome, metabolome, enzymes) in one dataset. Doing this for the CD patient subsets allowed us to assess whether predicting disease status using multi-omics outperformed single-omic sPLS-DA models. The control sPLS-DA model also was similarly predictive of CD status as the individual -omic sPLS-DA models (AUC = 0.959) (Figures 2m and 2n). However, even though the control model has comparable AUC with single-omic models, the concatenated control has higher balanced error rate (CD = 0.45) than single omic models (Table 1). This indicated that unstructured combining of multi-omics alone does not enhance predictive power.

**Table 1.**
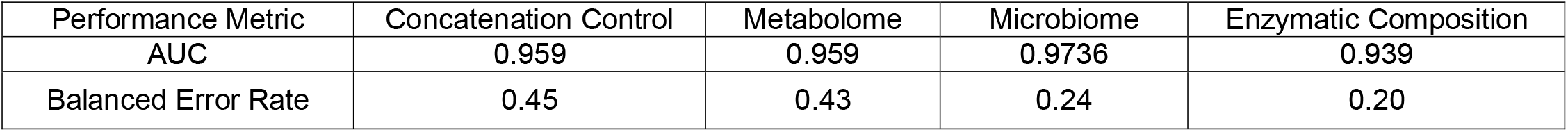
AUC and Balanced Error Rate comparison from single-omics sPLS-DA models for CD.

| Performance Metric | Concatenation Control | Metabolome | Microbiome | Enzymatic Composition |
| --- | --- | --- | --- | --- |
| AUC | 0.959 | 0.959 | 0.9736 | 0.939 |
| Balanced Error Rate | 0.45 | 0.43 | 0.24 | 0.20 |

sPLS-DA employs a regularization penalty when constructing LV’s, incorporating different features into different LV’s based on the ability to discriminate between Crohn’s Disease, and control status. The severity of this penalty is proportional to the number of features in the original dataset, making it much higher for the multi-omics sPLS-DA model compared to the single-omic sPLS-DA models. In our models for CD, 2 microbiome LV’s, explaining 5% and 2% variance encoded 100 and 2 features respectively, 3 metabolome LV’s explaining 17%, 3%, and 6% variance encoded 70, 40, and 90 features respectively, and 2 microbial enzyme LV’s, explaining 12% and 2% variance, encoded 100, and 2 features respectively.

By comparison, the concatenated data control sPLS-DA model resulted in lower feature coverage in the IBD microbiome multi-omics data compared to the single-omic sPLS-DA models. The control model that combines all omics for CD was comprised of 2 LV’s that explained 13% and 1% variance and encoded 70 features (0 microbes, 70 metabolites, 0 enzymes) and 1 feature (0 microbe, 1 metabolite, 0 enzymes) respectively. Given the number of features that encoded the covariance, individual microbiome, metabolome, and microbial enzyme sPLS-DA models show a higher combined data coverage than the control model (Table 2). It could be attributed to not restrictive Lasso penalties on individual models than the control model because of the small number of features in individual omics data sets than omics concatenation matrix data set.

**Table 2.** Single-omic sPLS-DA models show higher coverage than concatenation control model in CD.

| Latent Variable | Concatenation Control |  | Metabolome |  | Microbiome |  | Enzymatic Composition |  |
| --- | --- | --- | --- | --- | --- | --- | --- | --- |
|  | Encoded Features | Explained Variance | Encoded Features | Explained Variance | Encoded Features | Explained Variance | Encoded Features | Explained Variance |
| LV1 | 70 | 13% | 70 | 17% | 100 | 5% | 100 | 12% |
| LV2 | 1 | 3% | 40 | 3% | 2 | 2% | 2 | 2% |
| LV3 | 0 | 0% | 90 | 6% | 0 | 0% | 0 | 0% |
| Total | 71 | 16% | 200 | 20% | 102 | 7% | 102 | 14% |

### Structured integration of multi-omics latent variables Predicting Crohn’s Disease Status

We sought to test whether structured integration of multi-omics data better predicted disease status than single-omic sPLS-DA models or the concatenated control sPLS-DA models. Given AUC values and balanced error rates, we determine the single omic sPLS-DA models can classify control patients versus CD, and predictions are more accurate than concatenated control model. The most discriminative latent variables from these single sPLS-DA models were integrated in a structured multi-omics model. We first examined a main-effects only structured multi-omics model for predicting CD status. The form these models took was as a linear model with a slope intercept coefficient and main-effects coefficients for patient scores on each latent variable as features predicting disease status. A main-effects only regression model of microbiome, metabolite and microbial enzyme LVs was overall predictive of Crohn’s Disease status (p = 2.2 e-16). By evaluating the p-values from each data-type latent variables, we determine that the 2 Metabolomics LV’s emerge as the strongest data type to predict CD status from control patients in comparison with microbiota and enzymatic composition (Table 3).

**Table 3.** LIVE Modeling Outcome Table with Coefficients and p-values for CD.

| Crohn Disease Coefficient and p-values |  | Main Effects |  | Interactions |  |
| --- | --- | --- | --- | --- | --- |
| Feature |  | coefficients | p-value | coefficients | p-value |
| Metabolite LV1 |  | -0.0579 | < 2 e-16 | -0.0541 | 1.36 e-13 |
| Metabolite LV2 |  | 0.0624 | 3.98e-16 | 0.0572 | 8.10 e-08 |
| Metabolite LV3 |  | 0.0394 | 1.07e-10 | 0.0359 | 8.86 e-07 |
| Microbiota LV1 |  | 0.0256 | 0.141 | 0.0495 | 0.006 |
| Enzymes LV1 |  | 0.0115 | 0.083 | -0.0002 | 0.976 |
| Metabolite LV1 x Microbiota LV1 |  | - | - | 0.0079 | 0.007 |
| Metabolite LV2 x Microbiota LV1 |  | - | - | 0.0069 | 0.300 |
| Metabolite LV3 x Microbiota LV1 |  | - | - | -0.0049 | 0.264 |
| Metabolite LV1 x Enzymes LV1 |  | - | - | -0.0006 | 0.569 |
| Metabolite LV2 x Enzymes LV1 |  | - | - | 0.0078 | 0.006 |
| Metabolites LV3 x Enzymes LV1 |  | - | - | 0.0013 | 0.446 |
| Microbiota LV1 x Enzymes LV1 |  | - | - | -0.0048 | 0.118 |
| Metabolite LV1 x Microbiota LV1 x Enzymes LV1 |  | - | - | -0.0010 | 0.002 |
| Metabolite LV2 x Microbiota LV1 x Enzymes LV1 |  | - | - | 0.0007 | 0.176 |
| Metabolite LV3 x Microbiota LV1 x Enzymes LV1 |  | - | - | 0.00005 | 0.245 |
| Intercept |  | 0.6666 | < 2 e-16 | 0.7274 | < 2 e-16 |
|  |  | F-stat | 140.9 | F-stat | 77.84 |
|  |  | p-value | 2.2 e-16 | p-value | 2.2 e-16 |

We incorporated interaction effects between single-omic LV’s into the structured microbiome multi-omics PLS-PM framework to extract candidate microbe-metabolite-enzyme interactions driving CD. Incorporating microbiome, metabolome and microbial enzyme LV interaction effects into the Crohn’s Disease model identified four significant main effects from metabolome LV1, LV2 and LV3, as well as microbiota LV1. The model also identified significant interaction effects between microbiome LV1 - metabolome LV1, and metabolite LV2 - microbial enzyme LV1. The strongest microbe-metabolite-microbial enzyme LV interaction was present between metabolite LV1 - microbiota LV1 - enzymes LV1 which together separated CD patients from controls (Figure 3). This indicated that omic features allocated in these microbe-metabolite-microbial enzyme interacting Latent variables depends on the value of other omic variables to precisely define Crohn Disease status. In other words, a metabolite in conjunction with specific microbial species and microbial enzymes can determine CD status.

**Figure 3.**
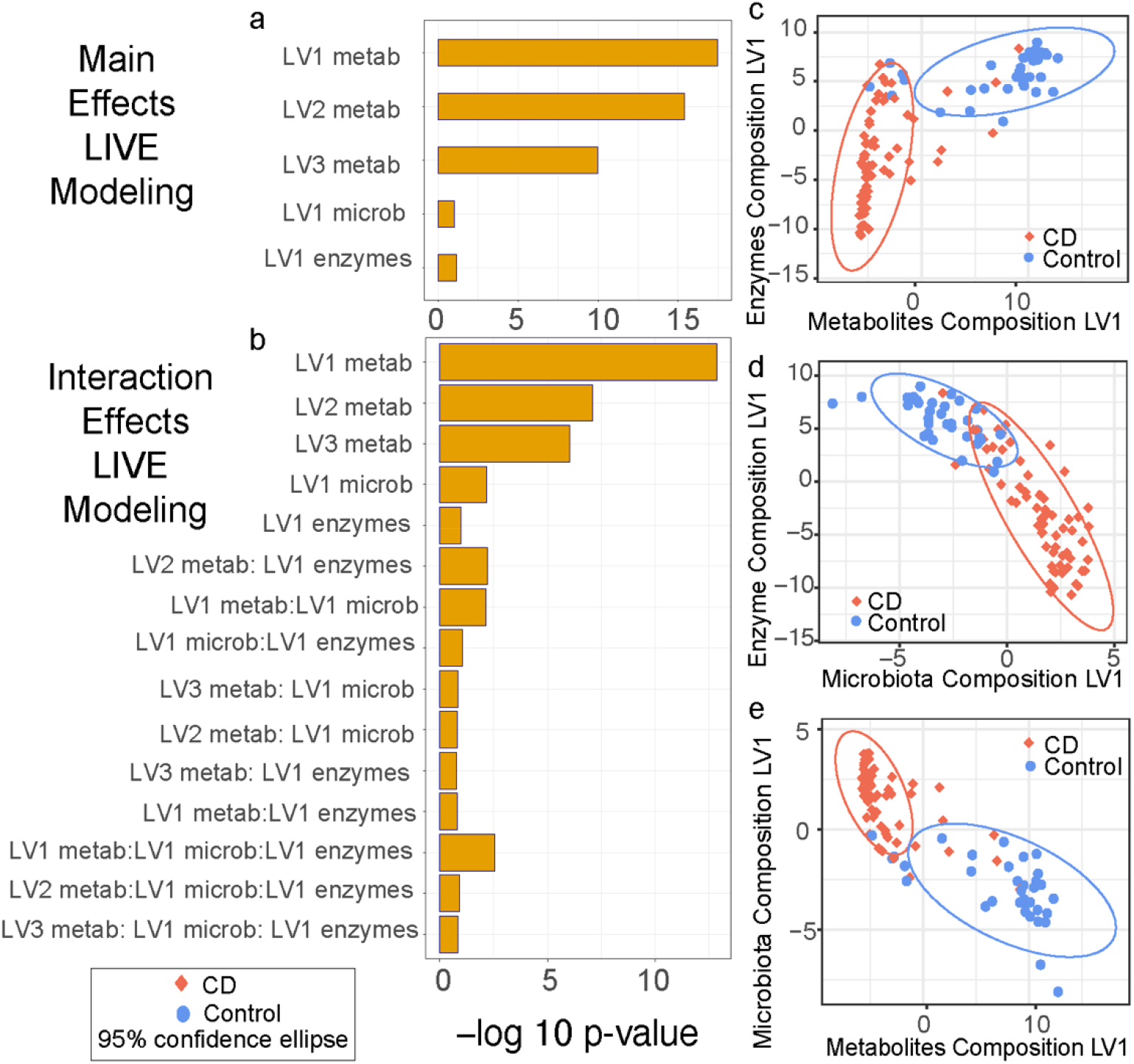
Main and Interaction Effects in a structured model regression for CD. (a) LV main effects model p-values. (b) LV Interaction effects p-values. (c) Scores plot of samples on enzymes LV1 and metabolites LV1. (d) Scores plot of samples on enzymes LV1 and microbiota LV1. (e) Scores plot of samples on microbiota LV1 and metabolites LV1.

### LIVE Modeling Prioritizes Multi-omic Features Associated with Crohn’s Disease

Having demonstrated that a structured PLS-PM of microbiome multi-omics data outperforms single-omic and concatenated multi-omics sPLS-DA models, we sought to extract biological insights from the predictive, interacting LV’s in our CD model. The most discriminant bacteria, metabolites and microbial enzyme features that determine disease status were prioritized from interacting LV’s by using as criteria Lasso Penalization and the variable importance of projection (VIP) scores of feature loadings on LVs (VIP > 1). After combining these two selection steps, correlation analysis was applied to identify microbe-metabolite-enzymes triads that were synergistically predictive of Crohn’s Disease status. The correlational analysis at this stage, restricted to features prioritized in our PLS-PM model, was prioritized to 23,188 microbe-metabolites-microbial enzymes pairs in the CD data based on the Lasso and VIP criteria. This represents a substantial decrease in the multiple testing burden of the original datasets of over 1.62 × 10^9^ microbe-metabolites-microbial enzymes pairs.

For Crohn’s Disease, analysis of these 23,188 microbe-metabolites-microbial enzymes pairs, identified 7,562 significant correlations (FDR q < 0.01) predictive of CD status on interacting multi-omic LV’s (FDR q <0.001 = 6,471 significant correlations; and FDR q < 0.0001 = 5305 significant correlations). LIVE modeling framework reduces the total number of features to a tractable number (7,562 CD) and the number of pairwise comparisons between features to that equivalent to a standard RNA-seq experiment. The prioritization of pairwise relationships between microbes, metabolites, and enzyme can be further refined by implementing more stringent thresholds on correlational coefficients, log fold change values, VIP scores, and correlation p-values or by framing interpretation in terms of a particular data type. We can visualize the impact of such thresholding by first constructing a microbe-metabolite-enzyme correlation network in Cytoscape [20] and then creating subnetworks that meet different thresholding criteria.

Here, we examine the effect of refining the significant pairwise correlations (i.e. pairwise local-structure) by increasing the threshold on the VIP score (i.e. global model importance) a given microbe, metabolite, or enzyme needs to achieve. For the CD model, we gradually increased the VIP threshold by 1, creating subnetworks of features with minimum global importance in our PLS-PM model until we reached a threshold that eliminated all bacterial nodes from the correlation network. At VIP score greater than 5 for Crohn’s Disease features, the network contained 72 nodes that include 44 metabolites, 27 microbial enzymes, and 1 bacterium, with a VIP threshold greater than 6 removing all bacteria. At our identified VIP threshold of 5, our model predicted *Coprococcus cactus* as one of the most discriminating bacterial features between CD and control samples, and Sphingolipids and Leucyl aminopeptidase as upregulated determinant metabolite and microbial enzyme features in Crohn’s Disease (Figure 4).

**Figure 4.**
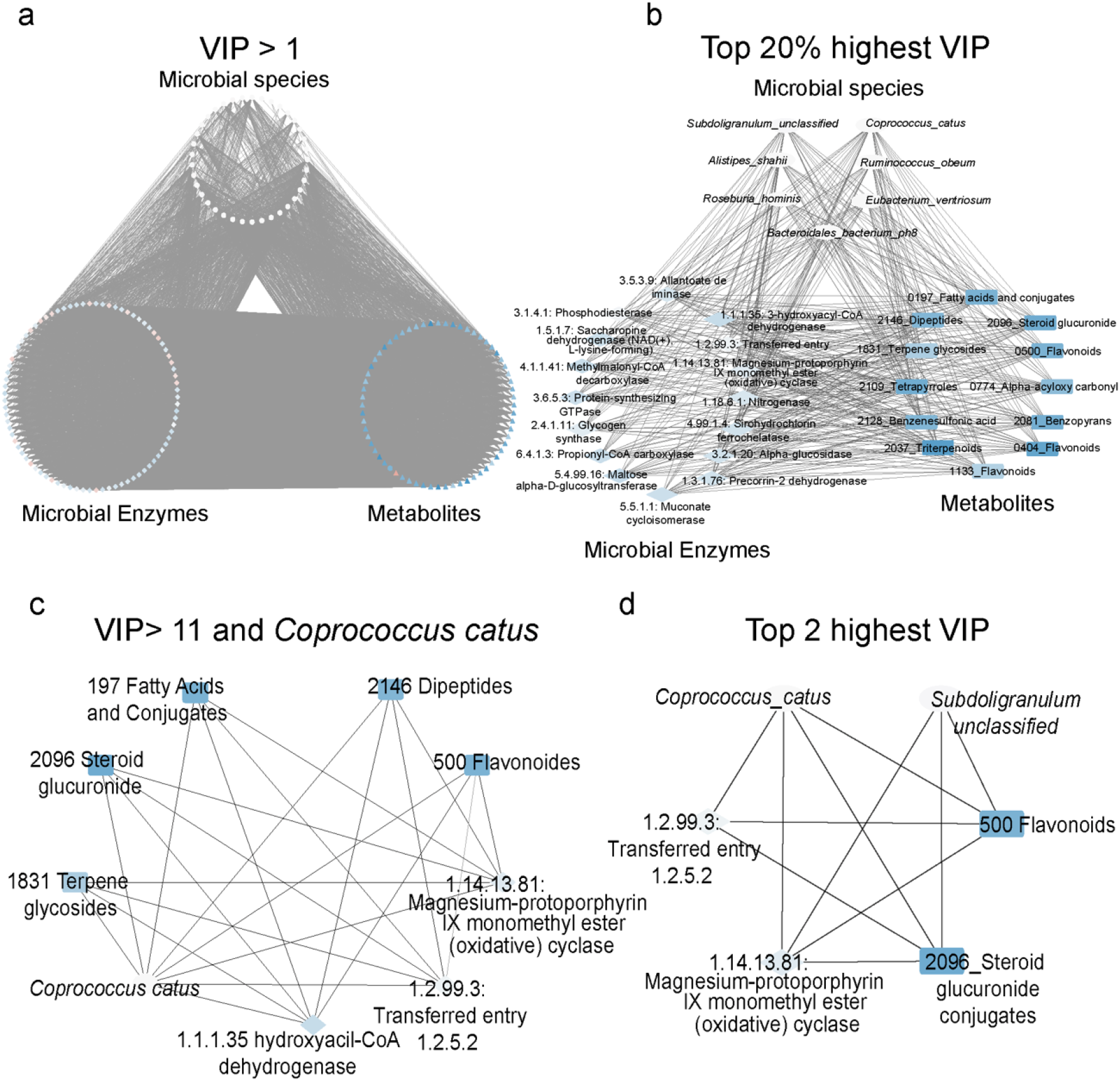
Meta Model Networks and Subnetworks for CD in Cytoscape. (a) Meta-Model networks VIP > 1. (b) Top 20% highest VIP per each omic data type. (c) VIP > 11 and *Coprococcus catus*. (d) Top 2 highest VIP. Edges reflect Spearman correlation coefficient.

### Single-omic sPLS-DA are as predictive of Ulcerative Colitis status as naive multi-omics models

For UC, the individual microbiome, metabolome, and microbial enzyme sPLS-DA models were significantly predictive of UC status (metabolome AUC = 0.932 Figure 5a and 5b, microbiome AUC = 0.969: Figure 5e and 5f, enzymatic composition AUC = 0.922: Figure 5i and 5j). Following LIVE modeling workflow, the most discriminative features of UC status were sorted by the PLS Variable Importance of Projection (Figure 5c, 5g, 5k) and the top features per each single sPLS-DA model were identified. The most discriminative metabolite features for UC are the diphenylmethanes, diterpenoids, pyridoxamines, tetrapyrroles and derivatives, cardiolipins, pyridinecarboxilic acids and monoalkyglycerophosphates (Figure 5d). The most relevant microbial species to classify UC patients are: *Gordonibacter pamelaeae, Roseburia hominis, Eubacterium rectale, Eubacterium hallii, Eubacterium ramulus, Alistipes shahii, Subdoligranulum unclassified*. (Figure 5h). The most discriminative enzymatic features of UC status are 2.7.1.66: Undecaprenol kinase, 1.5.3.1: Sarcosine oxidase, 2.7.1.66: Undecaprenol kinase, 1.5.3.1: Sarcosine oxidase, 2.4.1.4: Amylosucrase, 3.1.2.14: Oleoyl-[acyl-carrier-protein] hydrolase, 4.2.1.52: Transferred entry: 4.3.3.7, and 1.1.1.31: 3-hydroxyisobutyrate dehydrogenase (Figure 5l).

**Figure 5.**
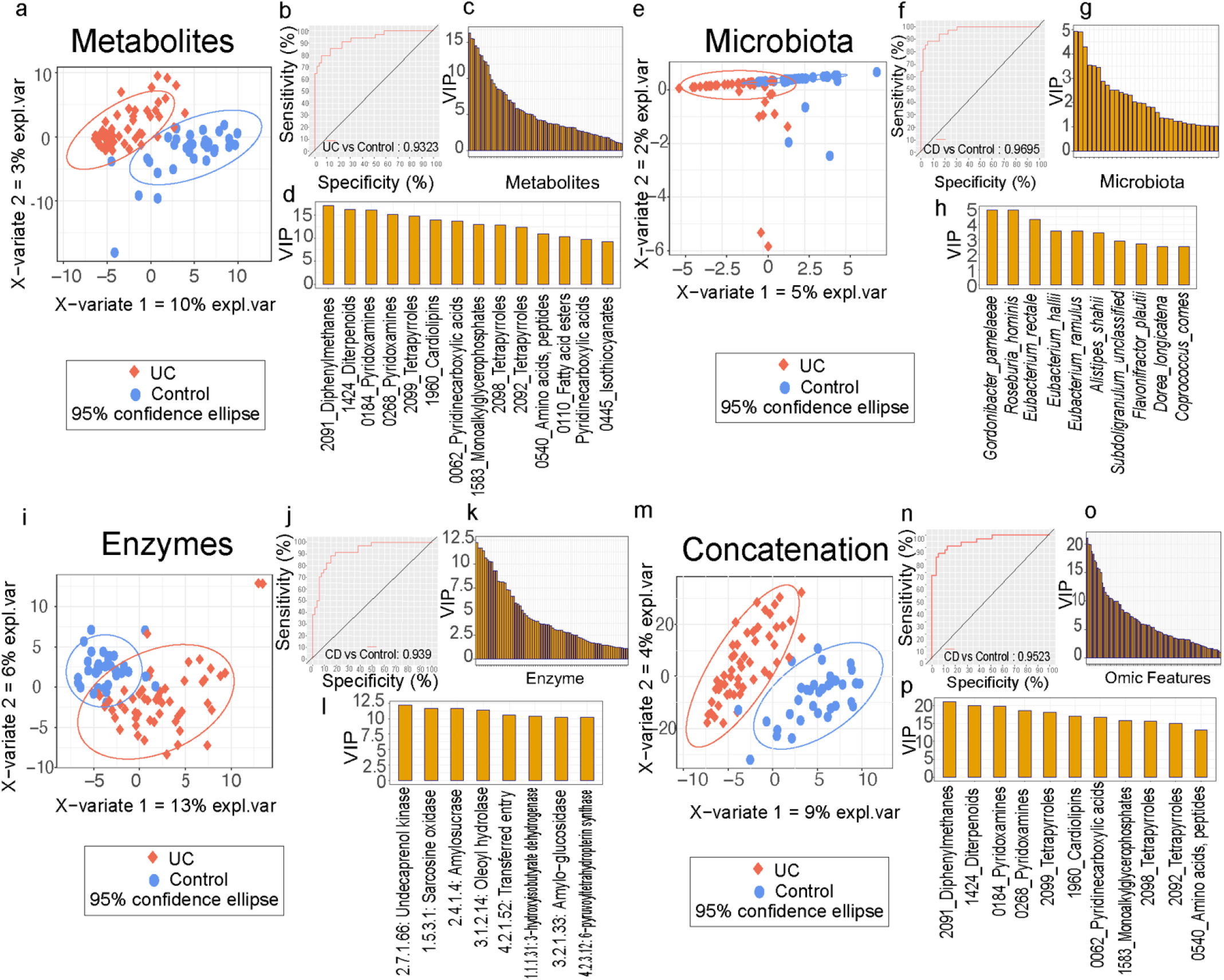
sPLS-DA model performance and variance importance of projections (VIP) for UC. (a) sPLS-DA sample scores plot for metabolomics data. (b) ROC curve for metabolomics sPLS-DA model. (c) VIP score histogram for metabolites. (d) Top 14 metabolites by VIP score. (e) sPLS-DA sample scores plot for microbiota composition data. (f) ROC curve for microbiota sPLS-DA model. (g) VIP scores histogram for microbiota. (h) Top 10 microbial taxa VIP scores. (i) sPLS-DA sample scores plot for enzymes data. (j) ROC curve for enzymes sPLS-DA model. (k) VIP score histogram for enzymes. (l) Top 8 enzymes by VIP score. (m) sPLS-DA sample scores plot for concatenation control multi-omic model. (n) ROC curve for concatenation control model. VIP score histogram for concatenation control model. (p) Top 11 feature VIP scores for the control model.

By comparing single-omics sPLS-DA models with the concatenated matrix as a computational control, we determined for UC status, similarly to the CD study case, that the control model has comparable AUC with single-omic models (AUC = 0.959) (Figures 5m and 5n), but, has higher balanced error rate (UC = 0.5) than single omic models (Table 4). This supports that unstructured combining of multi-omics alone does not necessarily enhance predictive power.

**Table 4.** AUC and Balanced Error Rate comparison from single-omics sPLS-DA models for UC.

| Performance Metric | Concatenation Control | Metabolome | Microbiome | Enzymatic Composition |
| --- | --- | --- | --- | --- |
| AUC | 0.952 | 0.922 | 0.932 | 0.969 |
| Balanced Error Rate | 0.5 | 0.343 | 0.5 | 0.125 |

The regularization penalty from sPLS-DA is proportional to the number of features in the original data set. Similar our CD study case, the penalty applied to the number of features for the multiomics sPLS-DA model is higher than single-omic sPLS-DA models, causing more error during predictions. For our UC sPLS-DA models, 2 microbiome LV’s, explaining 5% and 2% variance, encoded 80 and 1 features respectively, 2 metabolome LV’s, explaining 10%, and 3% variance, encoded 100 and 100 features respectively, and 2 microbial enzyme LV’s, explaining 13% and 6% variance, encoded 100, and 90 features respectively.

By comparison, the concatenated data control sPLS-DA model resulted in lower feature coverage in the UC microbiome multi-omics data compared to the single-omic sPLS-DA models. The control sPLS-DA model for predicting UC status was comprised of 2 LV’s, explaining 8% and 4% variance, and encoded 100 (0 microbes, 100 metabolites, 0 enzymes) and 1 feature (0 microbe, 1 metabolite, 0 enzymes) respectively (Table 5). Given the number of features that encoded the covariance, individual microbiome, metabolome, and microbial enzyme sPLS-DA models for UC show a higher coverage than the control model. Like the CD analysis, it could be attributed to not restrictive Lasso penalties on individual models than the control model because of the small number of features in individual omics data sets than omics concatenation matrix data set.

**Table 5.** Single-omic sPLS-DA models show higher coverage than concatenation control model in UC.

| Latent Variable | Concatenation Control |  | Metabolome |  | Microbiome |  | Enzymatic Composition |  |
| --- | --- | --- | --- | --- | --- | --- | --- | --- |
|  | Encoded Features | Explained Variance | Encoded Features | Explained Variance | Encoded Features | Explained Variance | Encoded Features | Explained Variance |
| LV1 | 100 | 8% | 100 | 10% | 80 | 5% | 100 | 13% |
| LV2 | 1 | 4% | 100 | 3% | 1 | 2% | 90 | 6% |
| Total | 101 | 16% | 200 | 13% | 81 | 7% | 190 | 19% |

### Structured integration of multi-omics latent variables Predicting Ulcerative Colitis Status

Given that single omic sPLS-DA models can classify control patients versus UC, and predictions are more accurate than concatenated control model, the structured integration of multi-omics data was implemented similar CD case study. The most discriminative latent variables of UC status from these single sPLS-DA models were integrated in a structured multi-omics model. Following the steps in LIVE modeling workflow, we first examined a main-effects only multi-omics model for predicting UC status, and we determined that a main-effects only regression model of microbiome, metabolite and microbial enzyme LVs was overall predictive of Ulcerative Colitis status (p = 2.2 e-16). By evaluating the p-values from each data-type latent variables, we determine that the 2 Metabolomics LV’s emerge as the strongest data type to predict CD status from control patients in comparison with microbiota and enzymatic composition (Table 6).

**Table 6.**
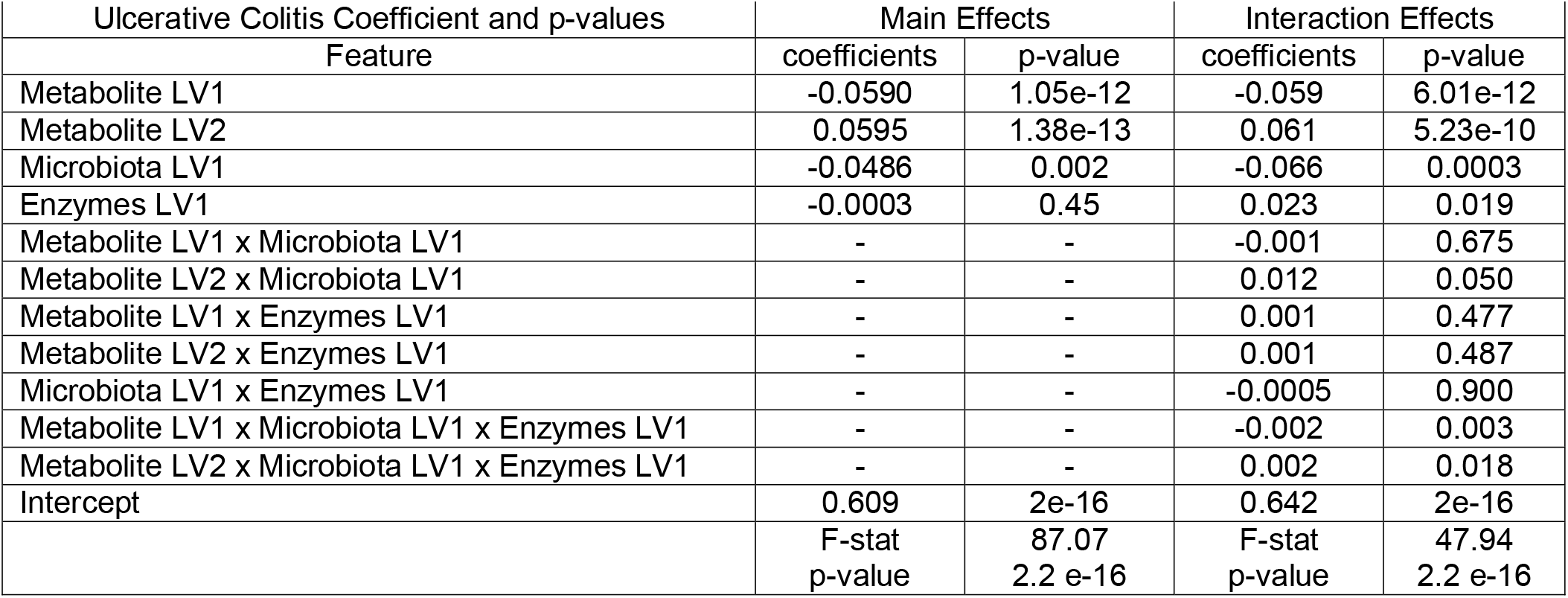
LIVE Modeling Outcome Table with Coefficients and p-values for UC.

We incorporated interaction effects between single-omic LV’s into the structured multi-omics model to extract candidate microbe-metabolite-enzyme interactions driving UC. Our LV’s interaction effects model for UC identified four significant main effects from metabolome LV1 and LV2, microbiota LV1, and microbial enzyme LV1. The model also identified significant interaction effects between metabolites LV2 - microbiota LV1, microbiome LV1 -metabolome LV2, and microbiome LV1 - microbial enzyme LV1. The strongest microbe-metabolite-microbial enzyme LV interaction was present between metabolite LV1 - microbiota LV1 - enzymes LV1 which together separated UC patients from controls (Figure 6). This indicated that the omic features projected in the strongest microbe-metabolite-microbial enzyme interacting Latent Variables depends on the value of other omic variables to precisely define Ulcerative Colitis status.

**Figure 6.**
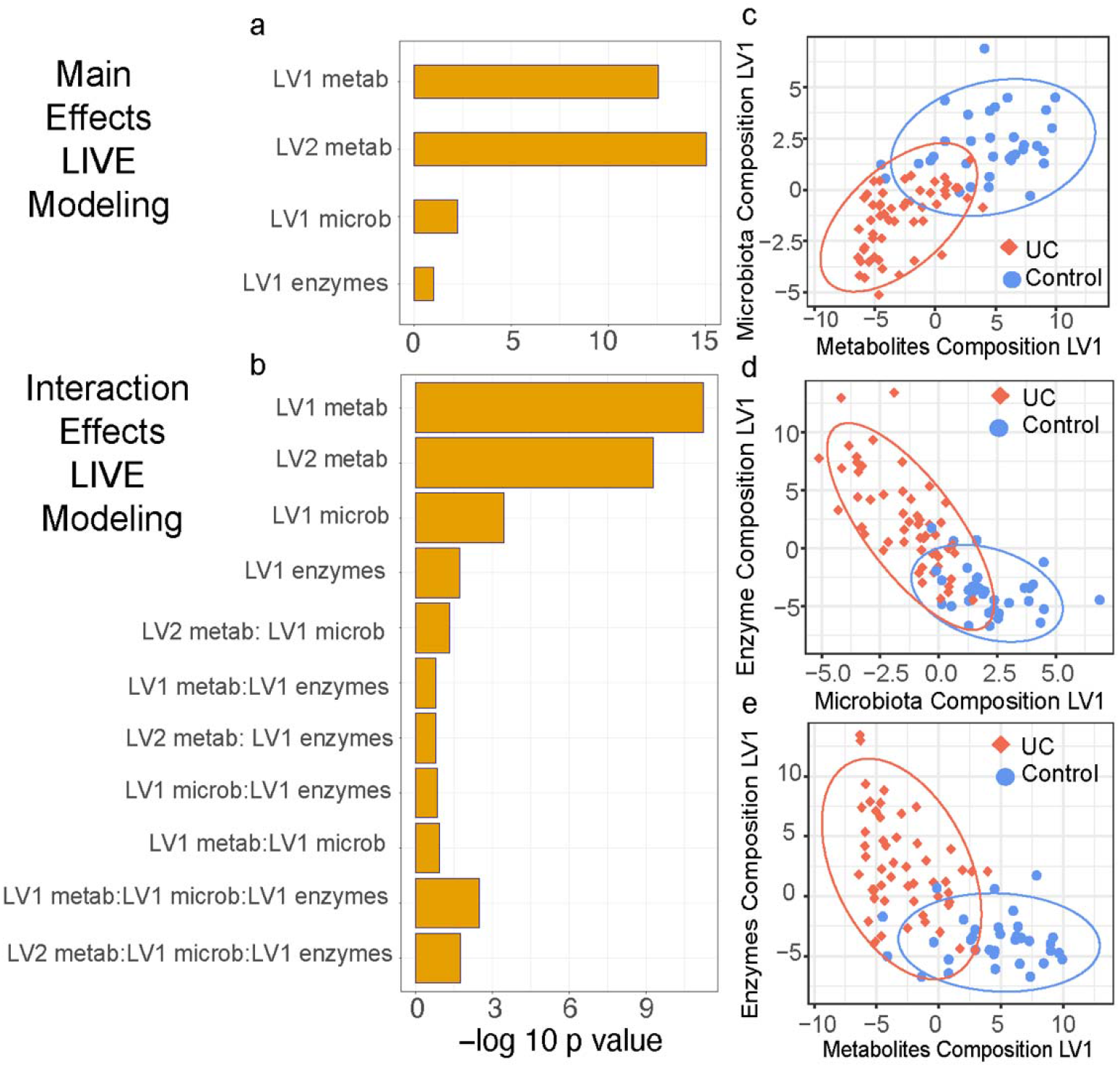
Main and Interaction Effects in a structured model regression for UC. (a) LV main effects model p-values. (b) LV Interaction effects p-values. (c) Scores plot of samples on microbiota LV1 and metabolites LV1. (d) Scores plot of samples on enzymes LV1 and microbiota LV1. (e) Scores plot of samples on enzyme LV1 and metabolites LV1.

### LIVE Modeling Prioritizes Multi-omic Features Associated with Ulcerative Colitis

The most discriminant bacteria, metabolites and enzyme features that determine Ulcerative Colitis disease status were prioritized from the predictive and interacting LV’s to extract biological insights, by using as criteria Lasso Penalization and the variable importance of projection (VIP) scores of feature loadings on LVs (VIP > 1). Like CD study case, by combining these two selection steps, and applying correlation analysis restricted to features prioritized in our PLS-PM model, the number of microbe-metabolites-microbial enzymes pairs for UC model was 21,514, a decrease in the feature space by 5 orders of magnitude from 1.62 × 10^9^ microbe-metabolites-microbial enzymes pairs in the original data. Analysis of these 21,514 microbe-metabolites-microbial enzymes pairs identified 2,271 significant correlations (FDR q < 0.01) (S5b Table: FDR q <0.001 = 935 significant pairs; FDR q < 0.0001 = 445 significant pairs) predictive of UC status on interacting LV’s.

In the microbe-metabolite-enzyme network for Ulcerative Colitis, VIP score was constrained greater than 4 and the model identified 69 nodes that include 39 metabolites, 27 microbial enzymes, and 3 bacteria, with a VIP score greater than 5 removing all bacterial features. Our UC model predicted *Eubacterium rectale, Gordonibacter parmelaeae* and *Roseburia hominis* as the most relevant bacterial features and showed several metabolites were specifically associated with each bacterium. For instance, the Tetrapyrroles and derivatives category is described as a downregulated metabolite associated with *Eubacterium rectale*. Fatty acid esters, Diphenylmethanes and Pyridoxamines as downregulated metabolites associated with *Gordonibacter pamelaeae*, and Tetrapyrroles and derivatives, Fatty acid esters, and Pyridoxamines as downregulated metabolites associated with *Roseburia hominis* and *Gordonibacter pamelaeae* in Ulcerative colitis (Figure 7).

**Figure 7.**
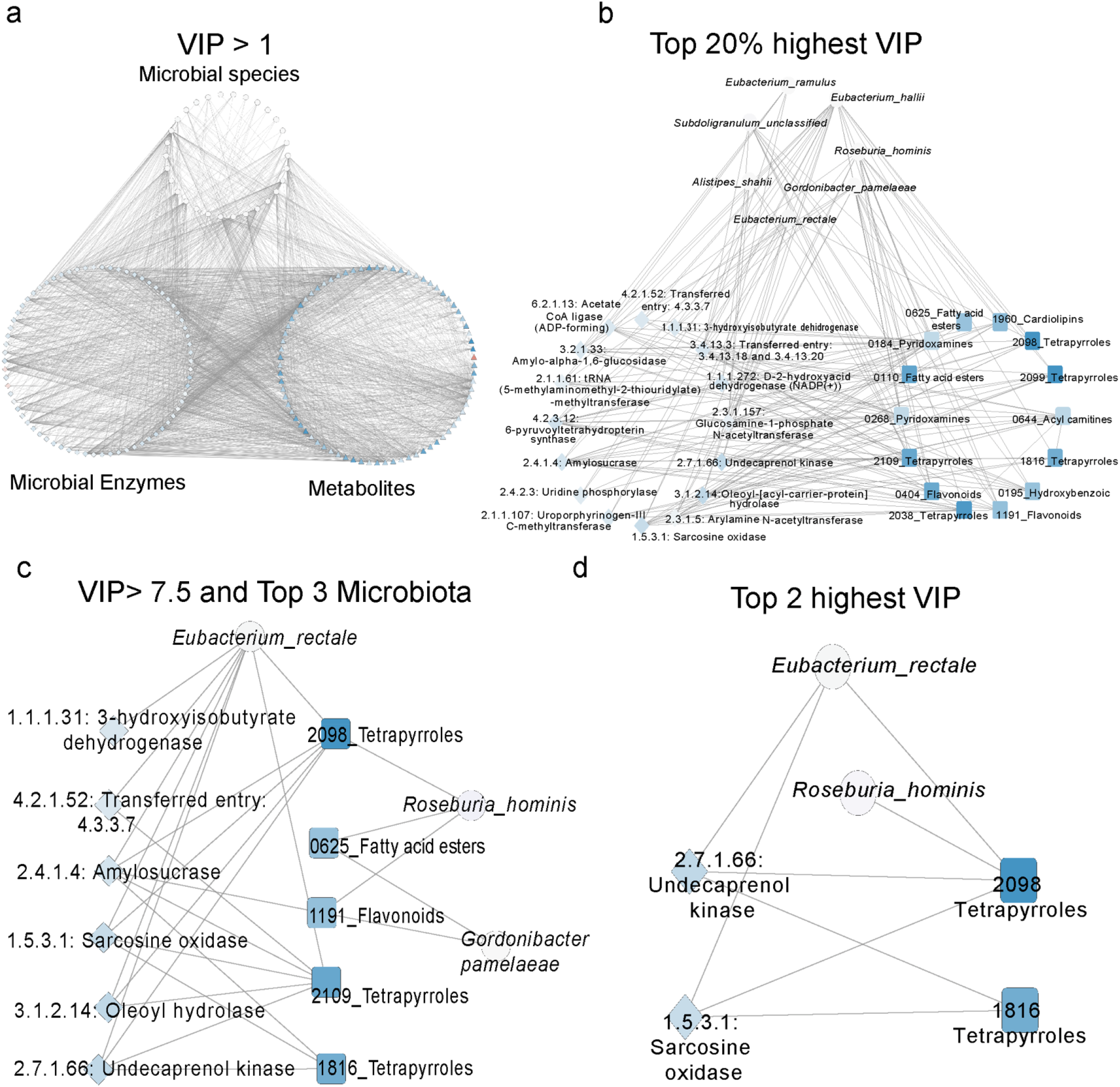
Meta Model Networks and Subnetworks for UC in Cytoscape. (a) Networks VIP > 1. (b) Top 20% highest VIP per each omic data type. (c) VIP > 7.5 and Top 3 microbial species. (d) Top 2 highest VIP. Edges reflect Spearman correlation coefficient.

## DISCUSSION

Our structural LIVE (Latent Interacting Variable Effects) modeling framework not only predicts features that are in concordance with previous studies, but also estimates novel omic associations that must be studied *in vitro* and *in vivo* models. This framework uncovers more feature interactions that increase the likelihood to establish testable hypothesis and studying biological mechanisms in IBD. LIVE modeling is a microbiome multi-omics PLS-PM framework reduces the total number of features to tractable numbers (7,562 CD and 2,271 UC FDR q < 0.01) and the number of pairwise comparisons between features to that equivalent to a standard RNA-seq experiment. This represents a substantial decrease in the multiple testing burdens of the original datasets. Additionally, given the greater number of significant correlations between CD microbiome multi-omic features than UC features reinforces the stratification of IBD into these two molecularly subtypes in addition to their differences in clinical presentation.

The prioritization of pairwise relationships between microbes, metabolites, and enzyme can be further refined in LIVE modeling by implementing more stringent thresholds on correlational coefficients, log fold change values, VIP scores, and correlation p-values or by framing interpretation in terms of a particular data type. We can visualize the impact of such thresholding by first constructing a microbe-metabolite-enzyme correlation network in Cytoscape and then creating subnetworks that meet different thresholding criteria. LIVE modeling workflow examines the effect of refining the significant pairwise correlations (i.e. pairwise local-structure) by increasing the threshold on the VIP score (i.e. global model importance) a given microbe, metabolite, or enzyme needs to achieve. It allows us to identify the most discriminative and interacting metabolites, microbial species and enzymes correlations for CD and UC.

This framework identified features associated with Inflammatory Bowel Diseases that have been reported previously by different studies [9-10]. Some of these features were reported by the original publication we drew data from, Franzosa *et al*. Here, the authors analyzed this data using other methods such as pairwise correlational analysis and random forest classifiers. In total, forty-four metabolites that were downregulated in Crohn’s Disease were predicted by our model at one of the highest levels of restriction (VIP > 5). Among the most important metabolite features are flavonoids, steroidal glucuronide conjugates, fatty acids and conjugates, dipeptides, terpene glycosides, tetrapyrroles and derivates, benzenesulfonic acids and derivatives, triterpenoids, benzopyrans and alpha-acyloxy carbonyl compounds [9, 21-23].

The Sphingolipids category was also predicted by our model as a determinant metabolite that was up regulated in Crohn’s Disease and reported by Franzosa *et al* and other studies [24]. From thirty-seven determinant microbiota species predicted by our model at the level of restriction of VIP >1 for Crohn’s Disease, thirteen were also reported by Franzosa *et al*. as representative microbial species. If the level of restriction is more stringent VIP>5, *Coprococcus catus* was predicted as the most determinant microbiota species and it also has been reported by Franzosa *et al*., but not prioritized as canonical feature, as well as in other studies [25-27]. Fold changes calculated from the relative abundance values of 37 predicted microbial species are about zero which is associated with the loss of species diversity, a characteristic phenomenon in IBD. From twenty-seven microbial enzymes predicted by our model at VIP > 5 for Crohn’s Disease, propionyl-CoA carboxylase, precorrin-2 dehydrogenase, leucyl aminopeptidase, NAD (+) synthase (glutamine-hydrolyzing) enzymes were also reported by Franzosa *et al*. as representative microbial enzymes.

Though bile acids and dicarboxylic acids were reported by Franzosa *et al*., these features were not predicted by our model at three distinctive levels of restrictions: VIP >5 (72 features), VIP > 1 (185 features), and q < 0.01 (7,562 significant features). We determined that these features were excluded in the single omic sPLS-DA models when individual omics classify between CD and control patients, it means that bile acids and dicarboxylic acids were not the most discriminative metabolites features of CD status. Several studies associated Bile acids with Inflammation chronic diarrhea which is connected to bile acid malabsorption in CD patients.

Our workflow generates several inferential statistics that facilitate prioritization of biological associations between microbes, metabolites, enzymes, and IBD status. In the preset IBD dataset, by using VIP, metabolite, microbiota, and enzyme interactions synergistically predict disease status can be visualized at different levels of restriction. These statistics provide flexibility in the to be more stringent with feature selection. For instance, in the Crohn’s Disease model, one of the highest levels of restriction was VIP > 5, and it estimated *Coprococcus catus* as the most determinant microbial species correlated with 44 metabolites and 27 microbial enzymes. Here, *Coprococcus catus* becomes the central focus of a network of metabolite and microbial enzyme interactions in Crohn’s Disease. Nevertheless, this framework can be even more selective if it is needed. If a higher resolution of feature interactions between metabolites and microbial enzymes with *Coprococcus catus* is required, the level of restriction can be incremented in the VIP regarding metabolites and microbial enzyme features. At VIP >11, our model estimate *Coprococcus catus* interact with 1.1.1.35: 3-hydroxyacyl-CoA dehydrogenase, 1.14.13.81: Magnesium-protoporphyrin IX monomethyl ester (oxidative) cyclase, and 1.2.99.3: aldehyde dehydrogenase; and the following metabolites: triterpenoids, benzenesulfonic acids and derivatives, tetrapyrroles and derivatives, terpene glycosides, dipeptides, fatty acids and conjugates, steroid glucuronide conjugates, and flavonoids (Figure 4c). This structural omic interaction framework that keeps the correlation structure within metabolites, microbiota, metabolome, and enzymes provides high-resolution property to evaluate omic interactions based on research needs.

A potential application of the LIVE Modeling framework’s identification of disease-driving molecules and taxa is to leverage features that discriminate disease status from control to identify biomarkers for drug response, diagnosis, and therapeutics in IBD. In 2021, Lee *et al*. reported features associated with anti-cytokine and anti-integrin therapies in this same cohort we analyzed in this study [11].. We compared the reported drug response features in Lee *et al*. with disease associated features selected by our structural interaction model. *Gordonibacter parmelaeae* was the main microbial species they identified as a determinant in drug response for Crohn’s Disease and Ulcerative Colitis, and it also was predicted by our structural correlational model for CD (VIP>1) and UC (VIP>4). Particularly in Crohn’s Disease, common microbial enzyme features predicted by our model and reported as drug response associated features were precorrin-4 C (11)-methyltransferase, precorrin-6B C (5,15) -methyltransferase (decarboxylating), glucosamine-1-phosphate N-acetyltransferase, and histidinol-phosphatase. For Ulcerative Colitis, the metabolite acyl carnitines, and the microbial enzymes arabinogalactan endo-beta-1,4-galactanase, and phosphoglucomutase (alpha-D-glucose-1,6-bisphosphate-dependent) were features associated with therapy response and disease status classification (Table 7). In sum, there are several commonalities between omics features associated with disease status in LIVE modeling and secondary association with drug response in IBD [9,11].

**Table 7.**
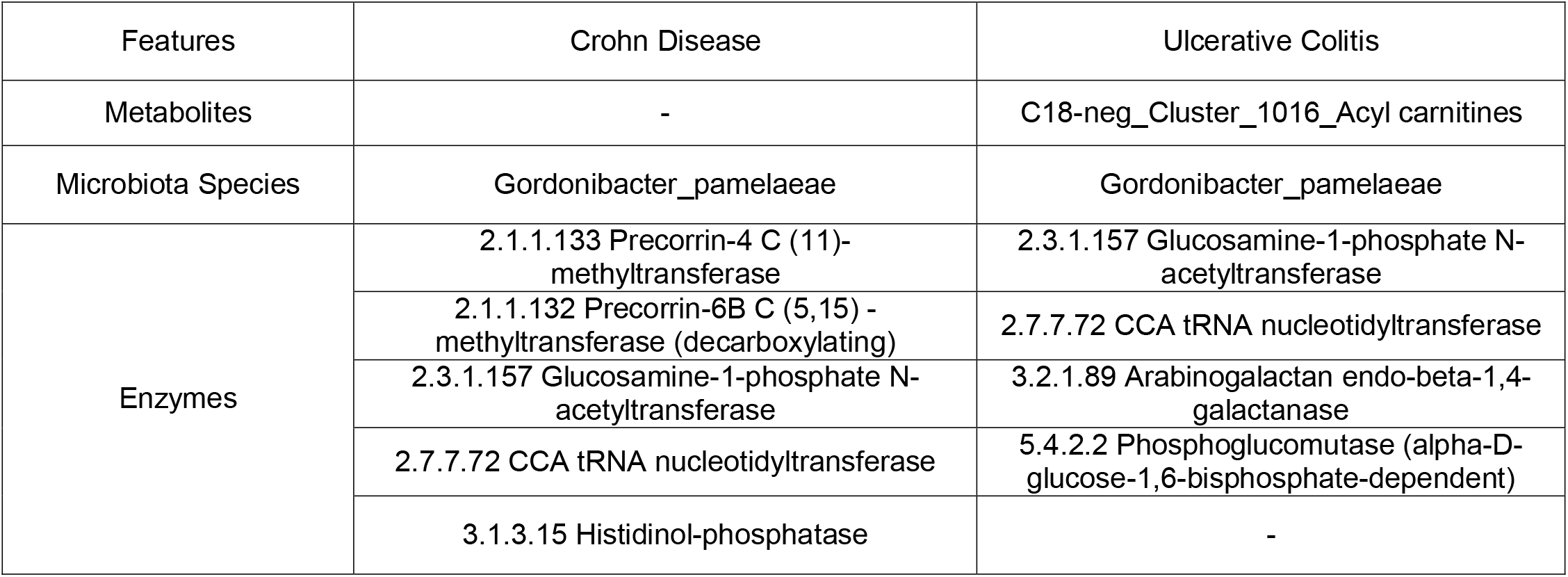
CD and UC features associated with drug response that were predicted by LIVE modeling.

LIVE modeling used the sample projections of the sPLS-DA model to build an interaction effects linear regression and predict omic features that are associated with disease and drug response in Inflammatory Bowel Diseases. These predictions were in good agreement with previous studies that reported disease and drug response features, which applying other statistical integrative methods. For instance, a combination of multivariable linear models and Random Forest classifiers with feature selection were applied to find associations between metabolome and microbiome features with clinical outcomes before and after exposure and integrate multi-omics and predict disease status and treatment response, respectively. Likewise, Metwaly *et al*. described linkages between IBD and sulfur metabolism using a Generalized Canonical Correlation Analysis (DIABLO-MixOmics) to integrated microbiota and metabolite profiles [24]. This last approach uses another correlation structure established by the weighted linear composites which vary to maximize the correlation between variables by overlapping their distributions [28]. Unlike LIVE Modeling however, it does not consider a regression modeling between sample projections of independent variables, and dependent variables.

Our LIVE Modeling framework makes a distinct and complementary contribution to these integrative methods that aim to predict metabolite and microbial species features that can predict IBD status. What distinguishes our approach is the flexibility to adapt to different types of microbiome multi-omics data, scalability for large and small cohort studies via reliance on latent variables and dimensionality reduction, and the intuitive interpretability of the linear meta-model integrating -omic data types. The results of LIVE modeling and the biological relationships can be represented in networks that connect local correlation structure of single omic data types with global community and omic structure in the latent variable VIP scores. This model arises as novel tool that allows researchers to be more selective about omic feature interaction without disrupting the structural correlation framework provided by sPLS-DA interaction effects modeling. It will lead to form testable hypothesis by identifying potential and unique interactions between metabolome and microbiome that must be considered for future studies.

## MATERIALS AND METHODS

### Preprocessing the Data

Publicly available metabolomics, and taxonomic and functional metagenomic data was obtained from 155 patients diagnosed with Crohn Disease, Ulcerative Colitis and non IBD patients (control) [9].. Relative Abundance profiles were log-transformed to variance stabilize the data with pseudo count 1 for zero values. This data set includes 201 microbiota features, 3829 metabolic features, and 2113 microbial enzyme features.

### Data Integration and Feature selection by sPLS-DA

We trained a sparse Partial Least Square – Discriminant Analysis (sPLS-DA) model on each single–omic data to predict the disease status using MixOmics R Package(17)(18). We used the function *tune. splsda* to select the optimal number of variables and components, and AUC plots were obtained with *auroc* function. Then, loadings, variance importance of projection (VIPs) and coefficients were exported from MixOmics.

### Multi-omics modeling with sample projection extracted from latent variables

Sample projections on the single-omic latent variables inferred from sPLS-DA models were used to train a generalized linear model with interaction effects terms. The alr4 R Package with the function lm were implemented to training the linear model. The main effects were patient projections on microbiome or metabolome, or microbial enzyme composition LV’s. Interactions effects were coded for each pair of microbiome, metabolome, and enzymatic LV’s. Significant interaction effects indicated that the predictive power of the bacterial loadings on a particular microbiome LV was conditioned upon the loadings of metabolites and enzymes on a particular metabolome and enzymatic LV.

### Feature Selection from Lasso Penalizations and Variance Importance Projection

From sPLS-DA models which identify the most predictive feature that classify disease status, we identified microbe, metabolite and enzymes features on significant interacting Latent variables with significant variable importance of projection (VIP) scores. The Lasso Penalization Formula is calculated by the following equation:

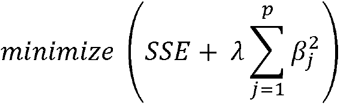

Variance Importance in Projection (VIP) scores for features are calculated using the regression coefficient *b*, weight vector *w*_*j*_, and score vector *t* as given in the following equation:

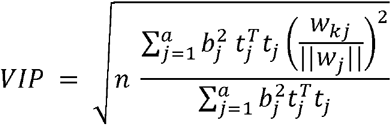

By considering Lasso Penalization and VIP greater than 1, we selected the most predictive and significant metabolic, microbiota and enzymes features that synergistically determine IBD status, for prioritized microbe-metabolite-enzyme correlation analysis. Correlational values were determined by Spearman correlational analysis. Duplicate correlational values were removed and same type omic data type interactions were removed. Then, p-value and false discovery rate were calculated from these omic matrices, and most relevant features were determined by using thresholds such as q-value < 0.01, and q-value < 0.001.

### Building Network Plots

Cytoscape software was used to visualize metabolites, microbiota, and enzyme interaction networks. To build these network plots, a node and edge files were used which containing the name and type of the features, log fold change and the VIP score, and correlation values, p value and false discovery rate, respectively. Different levels of restriction can be applied to visualize interaction network by filtering the data by VIP score, fold change, correlation, or other metadata.

## Acknowledgments

This work is based upon efforts supported by the EMBRIO Institute, contract #2120200, a National Science Foundation (NSF) Biology Integration Institute. The views and conclusions contained herein are those of the authors and should not be interpreted as representing the official policies, either expressed or implied, of NSF or the U.S. Government. The U.S. Government is authorized to reproduce and distribute reprints for governmental purposes notwithstanding any copyright annotation therein.

## SUPPORTING INFORMATION AND CAPTIONS

### List of Supplemental Materials

1. Supplemental Table 1.- Linn VIP from Metabolomic single-omic sPLS-DA model for CD
2. Supplemental Table 2.- Linn VIP from Microbiota composition single-omic sPLS-DA model for CD
3. Supplemental Table 3.- Linn VIP from Enzymatic composition single-omic sPLS-DA model for CD
4. Supplemental Table 4.- Model Performance single-omic sPLS-DA models and concatenation control.
5. Supplemental Table 5.- Significant metabolic, microbiota and enzyme pairs that determine CD status
6. Supplemental Table 6.- Node File for CD in Cytoscape
7. Supplemental Table 7.- Edge File for CD in Cytoscape
8. Supplemental Table 8.- Node File for CD VIP > 5 to build Meta-Model Networks
9. Supplemental Table 9.- Linn VIP from Metabolomic single-omic sPLS-DA model for UC
10. Supplemental Table 10.- Linn VIP from Microbiota composition single-omic sPLS-DA model for UC
11. Supplemental Table 11- Linn VIP from Enzymatic composition single-omic sPLS-DA model for UC
12. Supplemental Table 12.- Model Performance single-omic sPLS-DA models and concatenation control.
13. Supplemental Table 13.- Significant metabolic, microbiota and enzyme pairs that determine UC status
14. Supplemental Table 14.- Node File for UC in Cytoscape
15. Supplemental Table 15.- Edge File for CD in Cytoscape
16. Supplemental Table 16.- Node File for UC VIP > 4 to build Meta-Model Networks

## References

1. Ramos GP, Papadakis KA. Mechanisms of Disease: Inflammatory Bowel Diseases. Mayo Clin Proc. 2019;94(1):155–65.

2. de Mattos BRR, Garcia MPG, Nogueira JB, Paiatto LN, Albuquerque CG, Souza CL, et al. Inflammatory Bowel Disease: An Overview of Immune Mechanisms and Biological Treatments. Mediators Inflamm. 2015; 2015:1–11.

3. Schirmer M, Garner A, Vlamakis H, Xavier RJ. Microbial genes and pathways in inflammatory bowel disease. Nat Rev Microbiol. 2019;17(8):497–511.

4. Graham DB, Xavier RJ. Pathway paradigms revealed from the genetics of inflammatory bowel disease. Nature. 2020; 27;578(7796):527–39.

5. Schirmer M, Smeekens SP, Vlamakis H, Jaeger M, Oosting M, Franzosa EA, et al. Linking the Human Gut Microbiome to Inflammatory Cytokine Production Capacity. Cell. 2016 Nov;167(4):1125-1136.e8.

6. Glassner KL, Abraham BP, Quigley EMM. The microbiome and inflammatory bowel disease. J Allergy Clin Immunol. 2020;145(1):16–27.

7. Karpinska-Leydier, K., Amirthalingam, J., Alshowaikh, K., Iroshani Jayarathna, A., Salibindla, D., Paidi, G., and Ergin, H. E. Correlation Between the Gut Microbiome and Immunotherapy Response in Inflammatory Bowel Disease: A Systematic Review of the Literature. Cureus (2021), 13(8):e16808.

8. Plichta DR, Graham DB, Subramanian S, Xavier RJ. Therapeutic Opportunities in Inflammatory Bowel Disease: Mechanistic Dissection of Host-Microbiome Relationships. Cell. 2019;178(5):1041–56.

9. Franzosa EA, Sirota-Madi A, Avila-Pacheco J, Fornelos N, Haiser HJ, Reinker S, et al. Gut microbiome structure and metabolic activity in inflammatory bowel disease. Nat Microbiol. 2019 Feb;4(2):293–305.

10. Lloyd-Price J, Arze C, Ananthakrishnan AN, Schirmer M, Avila-Pacheco J, et al. Multi-omics of the gut microbial ecosystem in inflammatory bowel diseases. Nature. 2019. 569(7758):655–62.

11. Lee JWJ, Plichta D, Hogstrom L, Borren NZ, Lau H, Gregory SM, et al. Multi-omics reveal microbial determinants impacting responses to biologic therapies in inflammatory bowel disease. Cell Host Microbe. 2021, 29(8), 1294–1304.e4.

12. Lin E, Lane H-Y. Machine learning and systems genomics approaches for multi-omics data. Biomark Res. 2017;5(1):2.

13. Lee LC, Liong C-Y, Jemain AA. Partial least squares-discriminant analysis (PLS-DA) for classification of high-dimensional (HD) data: a review of contemporary practice strategies and knowledge gaps. The Analyst. 2018;143(15):3526–39.

14. Tenenhaus M, Vinzi VE, Chatelin Y-M, Lauro C. PLS path modeling. Comput Stat Data Anal. 2005 1;48(1):159–205.

15. Sanchez G. Sanchez, G. (2013) PLS Path Modeling with R, Trowchez Editions. Berkeley, 2013. Trowchez Editions. 2013. Available from: http://www.gastonsanchez.com/PLSPath Modeling with R.pdf

16. Lê Cao K-A, Martin PG, Robert-Granié C, Besse P. Sparse canonical methods for biological data integration: application to a cross-platform study. BMC Bioinformatics. 2009; 26;10(1):34.

17. Lê Cao K-A, Boitard S, Besse P. Sparse PLS discriminant analysis: biologically relevant feature selection and graphical displays for multiclass problems. BMC Bioinformatics. 2011; 22;12(1):253.

18. Rohart F, Gautier B, Singh A, Cao K-AL. mixOmics: an R package for ‘omics feature selection and multiple data integration. PloS Comput Biol 2017;13(11): e1005752.

19. Priya, S., Burns, M.B., Ward, T. et al. Identification of shared and disease-specific host gene–microbiome associations across human diseases using multi-omic integration. Nat Microbiol 7.2022; 780–795.

20. Shannon P. Cytoscape: A Software Environment for Integrated Models of Biomolecular Interaction Networks. Genome Res. 2003; 1;13(11):2498–504.

21. Vezza T, Rodríguez-Nogales A, Algieri F, Utrilla MP, Rodriguez-Cabezas ME, Galvez J. Flavonoids in Inflammatory Bowel Disease: A Review. Nutrients. 2016; 9;8(4):211.

22. Neurath MF. Host–microbiota interactions in inflammatory bowel disease. Nat Rev Gastroenterol Hepatol. 2020;17(2):76–7.

23. Huang X, Chen Z, Li M, Zhang Y, Xu S, Huang H, et al. Herbal pair Huangqin-Baishao: mechanisms underlying inflammatory bowel disease by combined system pharmacology and cell experiment approach. BMC Complement Med Ther. 2020;20(1):292.

24. Brown EM, Ke X, Hitchcock D, Jeanfavre S, Avila-Pacheco J, Nakata T, et al. Bacteroides-Derived Sphingolipids Are Critical for Maintaining Intestinal Homeostasis and Symbiosis. Cell Host Microbe. 2019;25(5):668-680.e7.

25. Metwaly A, Dunkel A, Waldschmitt N, Raj ACD, Lagkouvardos I, Corraliza AM, et al. Integrated microbiota and metabolite profiles link Crohn’s disease to sulfur metabolism. Nat Commun. 2020;11(1):4322.

26. Sankarasubramanian J, Ahmad R, Avuthu N, Singh AB, Guda C. Gut Microbiota and Metabolic Specificity in Ulcerative Colitis and Crohn’s Disease. Front Med. 2020; 7:606298.

27. van der Giessen J, Binyamin D, Belogolovski A, Frishman S, Tenenbaum-Gavish K, Hadar E, et al. Modulation of cytokine patterns and microbiome during pregnancy in IBD. Gut. 2020;69(3):473–86.

28. Sørensen M, Kanatsoulis CI, Sidiropoulos ND. Generalized Canonical Correlation Analysis: A Subspace Intersection Approach. IEEE Trans Signal Process. 2021; 69:2452–67.

